# Genomic prediction for commercial traits using univariate and multivariate approaches in Nile tilapia (*Oreochromis niloticus*)

**DOI:** 10.1101/725143

**Authors:** Rajesh Joshi, Anders Skaarud, Mayet de Vera, Alejandro Tola Alvarez, Jørgen Ødegård

## Abstract

**Background:** Over the past three decades, Nile tilapia industry has grown into a significant aquaculture industry spread over 120 tropical and sub-tropical countries around the world accounting for 7.4% of global aquaculture production in 2015. Across species, genomic selection has been shown to increase predictive ability and genetic gain, also extending into aquaculture. Hence, the aim of this paper is to compare the predictive abilities of pedigree- and genomic-based models in univariate and multivariate approaches, with the aim to utilize genomic selection in a Nile tilapia breeding program. A total of 1444 fish were genotyped (48,960 SNP loci) and phenotyped for body weight at harvest (BW), fillet weight (FW) and fillet yield (FY). The pedigree-based analysis utilized a deep pedigree, including 14 generations. Estimated breeding values (EBVs and GEBVs) were obtained with traditional pedigree-based (PBLUP) and genomic (GBLUP) models, using both univariate and multivariate approaches. Prediction accuracy and bias were evaluated using 5 replicates of 10-fold cross-validation with three different cross-validation approaches. Further, impact of these models and approaches on the genetic evaluation was assessed based on the ranking of the selection candidates.

**Results:** GBLUP univariate models were found to increase the prediction accuracy and reduce bias of prediction compared to other PBLUP and multivariate approaches. Relative to pedigree-based models, prediction accuracy increased by ∼20% for FY, >75% for FW and >43% for BW. GBLUP models caused major re-ranking of the selection candidates, with no significant difference in the ranking due to univariate or multivariate GBLUP approaches. The heritabilities using multivariate GBLUP models for BW, FW and FY were 0.19 ± 0.04, 0.17 ± 0.04 and 0.23 ± 0.04 respectively. BW showed very high genetic correlation with FW (0.96 ± 0.01) and a slightly negative genetic correlation with FY (−0.11 ± 0.15).

**Conclusion:** Predictive ability of genomic prediction models is substantially higher than for classical pedigree-based models. Genomic selection is therefore beneficial to the Nile tilapia breeding program, and it is recommended in routine genetic evaluations of commercial traits in the Nile tilapia breeding nucleus.

## Background

Over the past three decades, Nile tilapia (*Oreochromis niloticus*) industry has grown into a significant aquaculture industry spread over 120 tropical and sub-tropical countries around the world accounting for 7.4% of global production in 2015 [1]. Nile tilapia has also been called the “aquatic chicken” [2] as it is well-suited for aquaculture in wide range of trophic and ecological adaptations, from backyards to intensive cages. Since the early days, the industry has recognized the potential gains from selective breeding and the challenge was to develop a strain, suitable for production across varieties of production environments. This led to the establishment of the Genetically Improved Farmed Tilapia (GIFT) base strain in early 1990s by the crossing of 8 different Nile tilapia strains from Africa and Asia [3]. This GIFT strain has been widespread over the world and serves as the base in majority of the farmed Nile tilapia. GenoMar Supreme Tilapia (GST®) strain was derived from GIFT and has undergone 27 generations of selection for growth, fillet yield and robustness.

For a long time, the aquaculture breeding industry has relied on pedigree information for genetic improvement, but in the last half-decade, top international breeding companies have started to use routine genomic selection and other genomic technologies in their genetic improvement programs for Atlantic salmon [4], catfish [5], common carp [6] and rainbow trout [7]. Tilapia has two genome assemblies [8,9], five linkage maps of varying resolutions constructed using different types of markers [10–14] and two recent 50K SNP-Arrays [14,15]. With these recent developments in SNP-Arrays and HD linkage maps being supported by the commercial industries, it is believed that this has opened a new door of the genomic era in Nile tilapia also.

Genomic selection helps to utilise the within- and between-family variation in the population, even for sib-evaluated traits. For such traits, the pedigree-based classical selection methods are just able to utilise between-family variation [16]. Across species, including aquaculture, genomic selection methods has been shown to increase the predictive ability and genetic gain by deriving more accurate breeding values [17,18]. Hence, the first aim of this paper is to perform genetic analysis using either genomic and pedigree-based information in univariate and multivariate statistical models for the commercial traits in Nile tilapia. The second objective is to compare the predictive abilities of the pedigree- and genomic-based models.

## Methodology

### Experimental design and rearing procedure

The study was carried out on generation 26 of the GST® strain of Nile tilapia, which is a continuation of the GIFT program [3]. Each generation of GST® consists of 8 batches that follow a revolving breeding scheme where males from batch *n* are mated to females from batch *n-1.* This way only about 30 families are produced in each batch, significantly reducing the age difference within a batch compared to spawning all the 250 families in a generation at once. The families in one batch were created by mating the selected parents in a 1:1 mating design, where one male and one female were placed in a small breeding hapa. After mating, eggs were collected, and the families were kept separate until hatching.

After hatching, 40 fries were randomly selected from each family and pooled together, which were then reared in a nursery pond for 4 weeks and treated with hormones to produce an all-male population, mimicking the normal practice in commercial operations. After the nursery stage, they were then transferred to larger pond for a 30 week grow-out period.

### Harvesting

Fish from the experiment were grown for the entire 30-week period without any selection. At the end, all the surviving fish were slaughtered and measured for three commercial traits: body weight at harvest (BW), fillet weight (FW) and fillet yield (FY).

### Pedigree

True pedigree was unavailable, since all the offspring were reared communally immediately after hatching to reduce the maternal environmental and/or full-sib and/or tank effects. Thus, lateral fin clips were obtained for microsatellite parentage assignment and pedigree was constructed as described in [19]. This is the routine pedigree construction method in the commercial production of GST® strain and micro-satellite constructed pedigrees were available for the last 14 generations (i.e. pedigree back to generation 12 with the records of 110,900 fish). Since one male was mated to 1 female in each of the 253 families, only full-sibs were present in the dataset.

### Genotypes

Lateral fin clips were obtained for DNA extraction during harvesting. DNA extraction was done at BioBank (https://biobank.no/) and sent to CIGENE lab, NMBU (https://cigene.no/) for genotyping using Onil50® array [14]. The raw dataset contained 58,466 SNPs. Of these, 50,275 SNPs (86.75%) were classified as “PolyHighResolution” (formation of three distinctive clusters of homozygous and heterozygous genotype) and “NoMinorHom” (formation of two distinctive clusters with one homozygous genotype missing) using Axiom Analysis Suite Software [20]. These high-resolution genotypes were further cleaned for low minor allele frequency (MAF <0.05) using PLINKv1.07 [21] and the remaining 48,960 SNPs (83.74%) were used for genomic analysis. Similarly, 3 animals were filtered for low genotyping call rate (individual call rate <0.9) and only the 1444 animals with the phenotype, pedigree and genotypes were used for further statistical analysis. The final dataset contained 188 full-sib families with an average of 7.68 offspring per full-sib family (range 1 to 15; standard deviation = 4.48).

### Statistical analysis

Statistical analysis for three commercial traits was performed using two different approaches, namely univariate and multivariate, and two different models (PBLUP and GBLUP) within each approach; as described below

### Univariate approach

DMUv6 [22] was used to fit mixed linear models, using REML to estimate the variance components, heritability and the breeding values. Univariate BLUP models were used for the three commercial Nile tilapia traits described as;

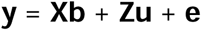

where, **y** is the vector of phenotypes, **b** is the vector of fixed effects that account for batch (7 levels), difference of age during harvesting (15 levels), filleter for the traits FW and FY (2 levels); **u** is the vector of random genetic effects; **e** is the vector of the residual errors; and **X** and **Z** are the corresponding design matrices for the fixed and random effects. For PBLUP, the distributional assumption of the random effects was multivariate normal, with mean zero and

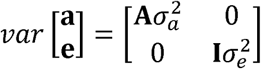

Where, 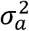 and 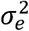 are additive genetic variances and residual variance respectively, **A** is the numerator relationship matrix obtained using micro-satellite generated pedigree and **I** is an identity matrix. The phenotypic variance was calculated as 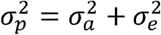 and the heritability (h^2^) was calculated as the ratio of 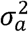 and 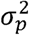.

For GBLUP, the numerator relationship matrix **A** was replaced with the genomic relationship matrix (**G**). The **G** matrix was constructed [23] as follows:

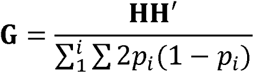

where **H** is a centered marker matrix, the sum in the denominator is over all loci and *p*_*i*_ is the allelic frequency at locus *i*.

### Multivariate approach

Multivariate models were built on the univariate models and are described as;

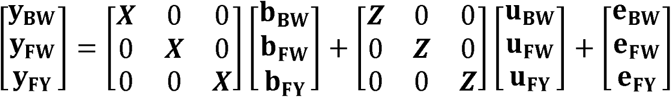

where, the symbols represent the same vectors as described in the univariate analysis, with the subscripts BW, FW and FY denoting the traits the vectors are associated with. For PBLUP models in multivariate approach, the distributional assumption of the random effects are structured as;

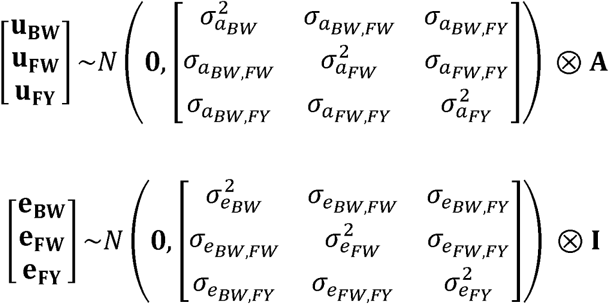

The symbols represent the same variance components as described in the univariate analysis with the subscripts denoting the trait the variance components are associated with. The elements 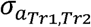 and 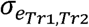 denotes the genetic and residual covariances between two traits, with the subscripts.

For GBLUP models, the numerator relationship matrix was replaced by the genomic relationship matrix (**G**).

### Predictive ability

Comparison between the predictive ability of PBLUP and GBLUP models was performed by both univariate and multivariate approaches using 5 replicates of 10-fold cross validation in different cross-validation methods. 10-fold cross validation allows us to mask the phenotypes of ∼10% of animals, which is predicted using the phenotypes of the rest of the 90% phenotypes.

Three different cross-validation methods were used to quantify the prediction accuracy of the models. With the “random cross-validation” method, the dataset was randomly divided into 10 batches, predicting one batch at a time using the phenotypes of the remaining 9 batches. Similarly, with “within family cross-validation” method, the phenotypes of (as close as possible to) 10% of the animals within a full-sib family are masked and phenotypes of the unmasked members of the family and other families are used to predict the masked phenotype. This scenario is important with the sib-testing strategy usually done for invasively measured traits like FY. Finally, with the “across family cross-validation” method, the phenotypes of all the animals in a full-sib are masked and the phenotypes of the individuals from other families are used to predict the masked phenotype. This scenario is appropriate where phenotype collection for all the population is very expensive and we measure the phenotypes in few families only or in different cohorts of fish.

Predictive ability of the GBLUP and PBLUP models were calculated as the Pearson’s correlation between GEBVs or EBVs of all predicted phenotypes adjusted for the fixed effects in one replicate. Results were averaged over the 5 replicates. The obtained mean value of correlation was converted to the expected prediction accuracy by dividing the correlation coefficient by the square root of the heritability. Heritabilities obtained from multivariate genomic models were used to assess the prediction accuracy. Standard error of prediction accuracy was calculated as [24];

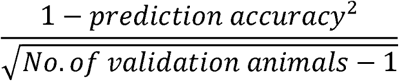

In addition, regression coefficient of phenotypes adjusted for the fixed effects on GEBVs or EBVs were used as to assess the bias of the prediction. Theoretically, a regression coefficient of 1 indicates unbiased prediction, whereas the value <1 indicates inflation of GEBV or EBV and >1 indicates deflation of GEBV or EBV. The mean value and standard error of the mean of the regression coefficient was calculated from the five replicates.

## Results

### Descriptive Statistics

Descriptive statistics for the three traits: BW, FW and FY are presented in Table 1. The mean (± standard deviation) phenotypic measurements for BW, FW and FY were 817.37 (± 261.11) g, 300.01(± 107.34) g and 36.40% (± 2.5%), respectively. The coefficient of variation ranged from about 7% for FY to as high as 36% for fillet weight. The scatterplot and phenotypic correlations between the traits are presented in Supplementary Figure S1.

**Table 1:** Descriptive statistics for the three commercial traits of Nile tilapia

|  | Units | Min | Max | Median | Mean | Mean (SE) | SD | CV% |
| --- | --- | --- | --- | --- | --- | --- | --- | --- |
| **BW** | g | 138.70 | 1893.70 | 780.30 | 817.37 | 6.87 | 261.11 | 31.95 |
| **FW** | g | 39.10 | 754.60 | 284.25 | 300.01 | 2.82 | 107.34 | 35.78 |
| **FY** | % | 20.83 | 46.64 | 36.56 | 36.40 | 0.07 | 2.50 | 6.90 |

**Note:** Min is the smallest phenotype, Max is the largest phenotype, SD is the standard deviation, SE is the standard error and CV is the coefficient of variation expressed as percentage. The traits BW represents body weight at harvest, FW represents fillet weight and FY represents fillet yield.

### Estimates of heritabilities

Estimates of variance components and heritabilities using univariate and multivariate approaches are presented in Table 2, whereas the genetic and phenotypic correlation between the traits obtained using multivariate approach is presented in Table 3. All the traits were found to have medium heritabilities. GBLUP models were found to give lower heritability estimates compared to PBLUP models in both univariate and multivariate approaches. Heritabilities using multivariate approach were slightly higher for the traits BW and FW, compared to univariate approach.

**Table 2:** Heritabilities and variance parameters for PBLUP and GBLUP models using univariate and multivariate approaches.

| Approaches | Traits | Model | $\sigma_a^2$ | $\sigma_e^2$ | $\sigma_p^2$ | $h^2$ | se | $\sigma_a^{2*}$ | $h^{2*}$ |
| --- | --- | --- | --- | --- | --- | --- | --- | --- | --- |
| Univariate | BW | PBLUP | 7131 | 25394 | 32525 | 0.22 | 0.06 | 7262 | 0.22 |
|  | BW | GBLUP | 5467 | 26742 | 32209 | 0.17 | 0.04 | 5437 | 0.17 |
|  | FW | PBLUP | 1230 | 4076 | 5306 | 0.23 | 0.06 | 1253 | 0.24 |
|  | FW | GBLUP | 842 | 4384 | 5226 | 0.16 | 0.04 | 837 | 0.16 |
|  | FY | PBLUP | 1.80 | 3.69 | 5.49 | 0.33 | 0.07 | 1.83 | 0.33 |
|  | FY | GBLUP | 1.21 | 4.13 | 5.34 | 0.23 | 0.04 | 1.21 | 0.23 |
| Multivariate | BW | PBLUP | 9279 | 24068 | 33348 | 0.28 | 0.06 | 9449 | 0.28 |
|  | BW | GBLUP | 6168 | 26366 | 32534 | 0.19 | 0.04 | 6134 | 0.19 |
|  | FW | PBLUP | 1488 | 3930 | 5419 | 0.27 | 0.06 | 1516 | 0.28 |
|  | FW | GBLUP | 899 | 4369 | 5268 | 0.17 | 0.04 | 894 | 0.17 |
|  | FY | PBLUP | 1.82 | 3.68 | 5.50 | 0.33 | 0.07 | 1.85 | 0.33 |
|  | FY | GBLUP | 1.26 | 4.11 | 5.37 | 0.23 | 0.04 | 1.25 | 0.23 |

**Note:** $\sigma_a^{2*}$ and $h^{2*}$ are the additive genetic variance and heritability corrected to the base generation as per [25]. The additive variance was multiplied by mean of corresponding diagonal relationship matrix – mean of the corresponding relationship matrix and the heritability was calculated based on this variance parameter. The
mean of the diagonal and whole matrix for **A** were 1.018322 and 0, and for **G** were 0.9951903 and 0.000689405 respectively.

**Table 3:**
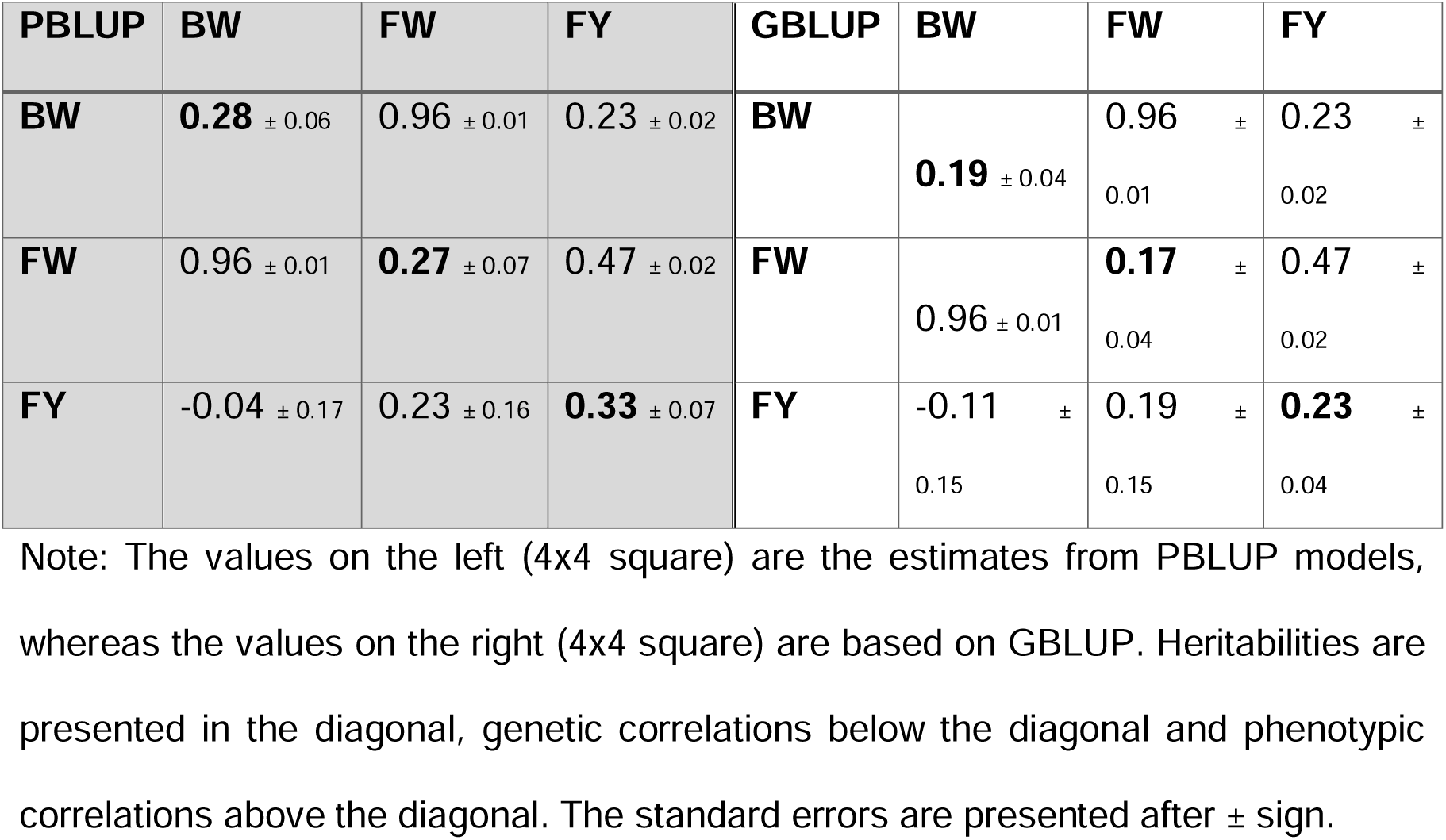
Heritabilities, phenotypic and genetic correlation using multivariate approach.

| PBLUP | BW | FW | FY | GBLUP | BW | FW | FY |
| --- | --- | --- | --- | --- | --- | --- | --- |
| **BW** | **0.28** ± 0.06 | 0.96 ± 0.01 | 0.23 ± 0.02 | **BW** | **0.19** ± 0.04 | 0.96 ± 0.01 | 0.23 ± 0.02 |
| **FW** | 0.96 ± 0.01 | **0.27** ± 0.07 | 0.47 ± 0.02 | **FW** | 0.96 ± 0.01 | **0.17** ± 0.04 | 0.47 ± 0.02 |
| **FY** | -0.04 ± 0.17 | 0.23 ± 0.16 | **0.33** ± 0.07 | **FY** | -0.11 ± 0.15 | 0.19 ± 0.15 | **0.23** ± 0.04 |

Note: The values on the left (4x4 square) are the estimates from PBLUP models, whereas the values on the right (4x4 square) are based on GBLUP. Heritabilities are presented in the diagonal, genetic correlations below the diagonal and phenotypic correlations above the diagonal. The standard errors are presented after ± sign.

The results indicated a slightly unfavorable genetic correlation between FY and BW (albeit non-significantly different from 0). The genetic correlations with the trait FY was higher for PBLUP models in multivariate approach, compared to GBLUP.

### Impact on the genetic evaluation

The correlation of the EBVs and/or GEBVs using two different approaches, namely univariate and multivariate, and two different models (PBLUP and GBLUP) within each approach are presented in Figure 2.

**Figure 1:**
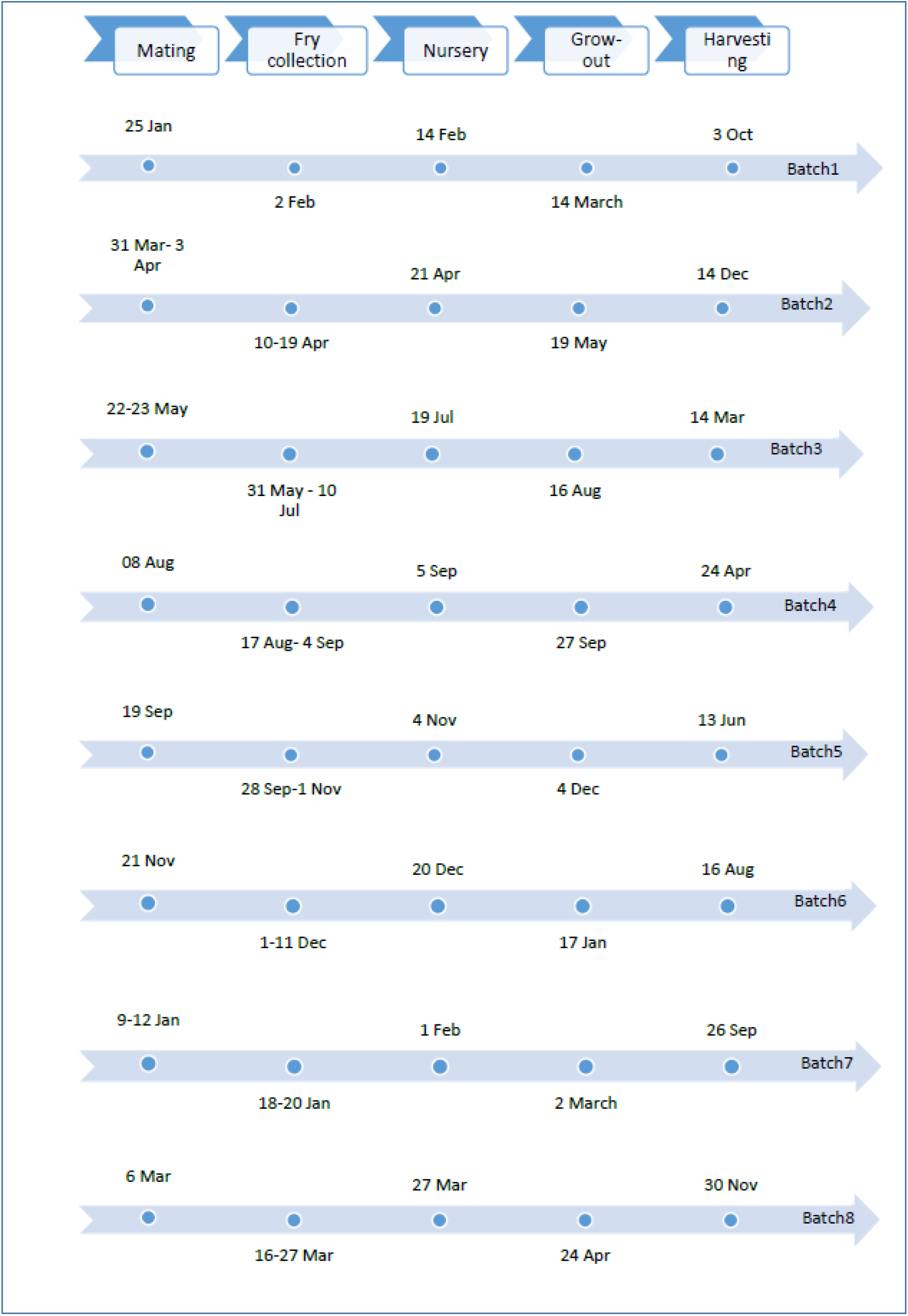
Dates showing different stages of lifecycle in Nile tilapia. The population were reared in 8 different batches during 2017-18

**Figure 2:**
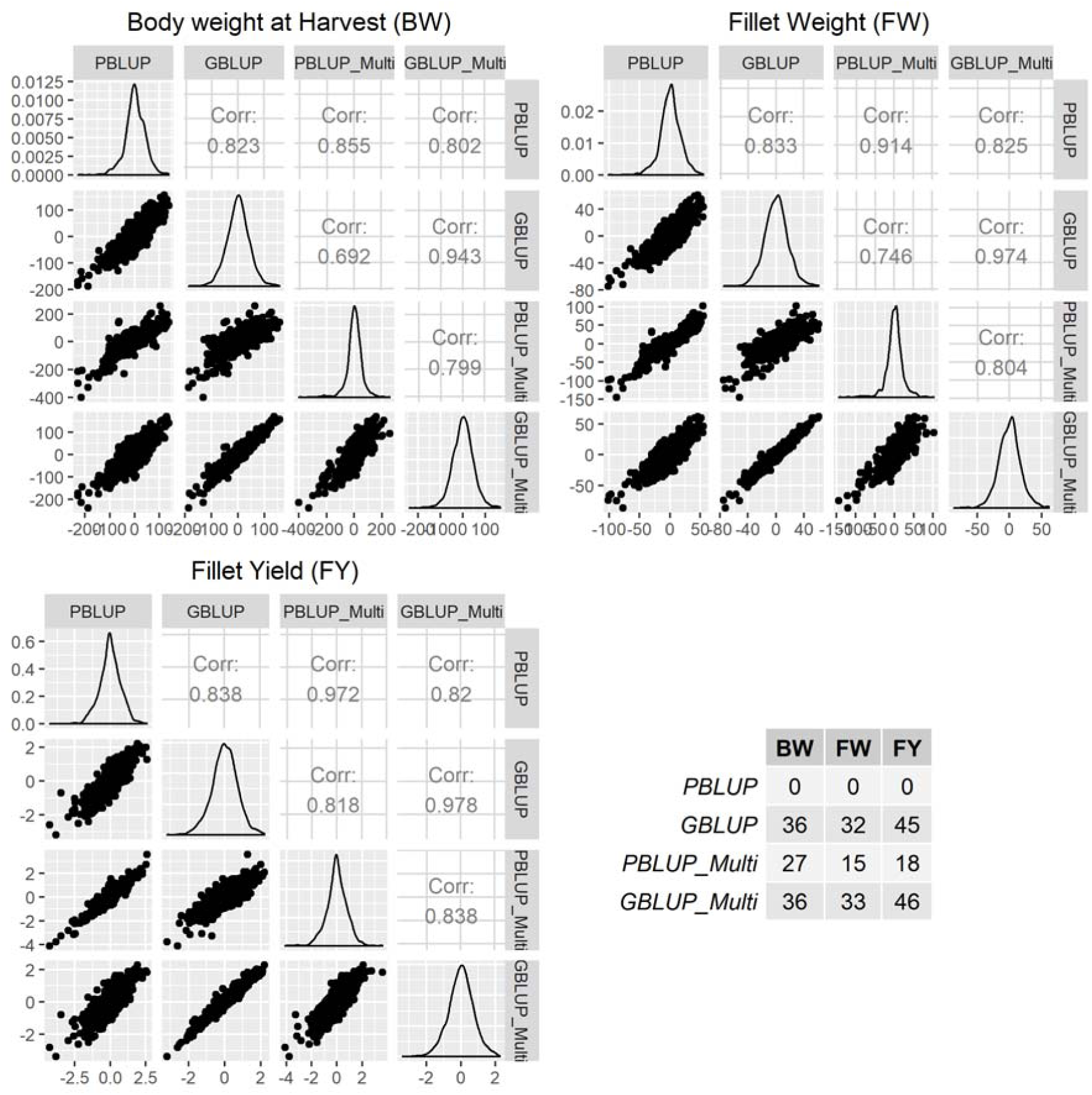
Impact of different models and approaches on the genetic evaluation. The models with univariate approach are shown as PBLUP and GBLUP, whereas the models with multivariate approaches have suffix “multi” in the models. The first three figures show the scatterplot and correlation between the EBVs and GEBVs for 3 different traits. The table on the bottom right axis shows the impact of model choice for the top 100 animals after ranking the animals based on EBVs or GEBVs. Since the comparison is based on PBLUP model in univariate approach, the 0 for PBLUP is by definition.

In general, the use of multivariate vs. univariate approaches affected the ranking of the breeding values, with correlations between EBVs/GEBVs ranging 0.86 to 0.98. There was less reranking among GBLUP univariate and multivariate approaches, compared with the PBLUP. Further, models within the same approach (i.e. PBLUP and GBLUP models within univariate and multivariate approaches) for three different traits revealed similar correlation in the range of 0.80 to 0.83. In overall, FY had higher correlation between models and approaches and BW had lowest correlation between models and approaches. Thus, FY showed the least differences and BW showed the major differences in the genetic evaluation by the use of different models and approaches, based on correlation of the EBVs. A lower correlation may indicate that careful selection of the model and approach has to be done, so that the genetic gain can be maximised.

These differences in the estimated breeding values also brought the change in the ranking of the 100 best animals (see table in the bottom right axis in Figure 2). Using PBLUP univariate approach as the reference group, major changes in the top 100 animals were observed using different models (PBLUP and GBLUP) and approaches (univariate and multivariate). GBLUP was less sensitive to univariate/multivariate modelling and the changes were more pronounced when going from PBLUP to GBLUP, which is consistent with the outcomes of the correlation of the breeding values. No major differences in the list of top 100 animals were observed using GBLUP univariate and GBLUP multivariate approaches, as these approaches also had the highest correlation of the estimated breeding values. These observations were similar across all the traits.

### Prediction accuracy

Estimates for the prediction accuracy in different cross-validation methods are presented in Figure 3. As expected, prediction accuracy was lower in “across-family” and similar in “random” and “within family” cross validation methods, for all three traits. Prediction accuracies using PBLUP models in across-family cross validation methods were found to be very low, while GBLUP models increased the prediction accuracy by 119% for FY to as high as 759% for BW. This huge increase in accuracy is expected, as the PBLUP models have very limited potential for across-family prediction in this material (no half-sibs available). For both random and within-family cross-validation methods, GBLUP models were found to increase the prediction accuracy by ∼20% for FY, >75% for FW and >43% for BW, compared to PBLUP models in univariate approach. Similar differences were found using PBLUP and GBLUP models in multivariate approach. In the majority of the cases (GBLUP and PBLUP), going from univariate to multivariate models did not improve prediction accuracy. However, for traits BW and FW in random cross-validation approach, a GBLUP multivariate model was found to slightly increase the prediction accuracy. In contrast, PBLUP multivariate models performed worse than univariate models, even giving negative prediction accuracy for BW and FY using the across-family cross validation method.

**Figure 3:**
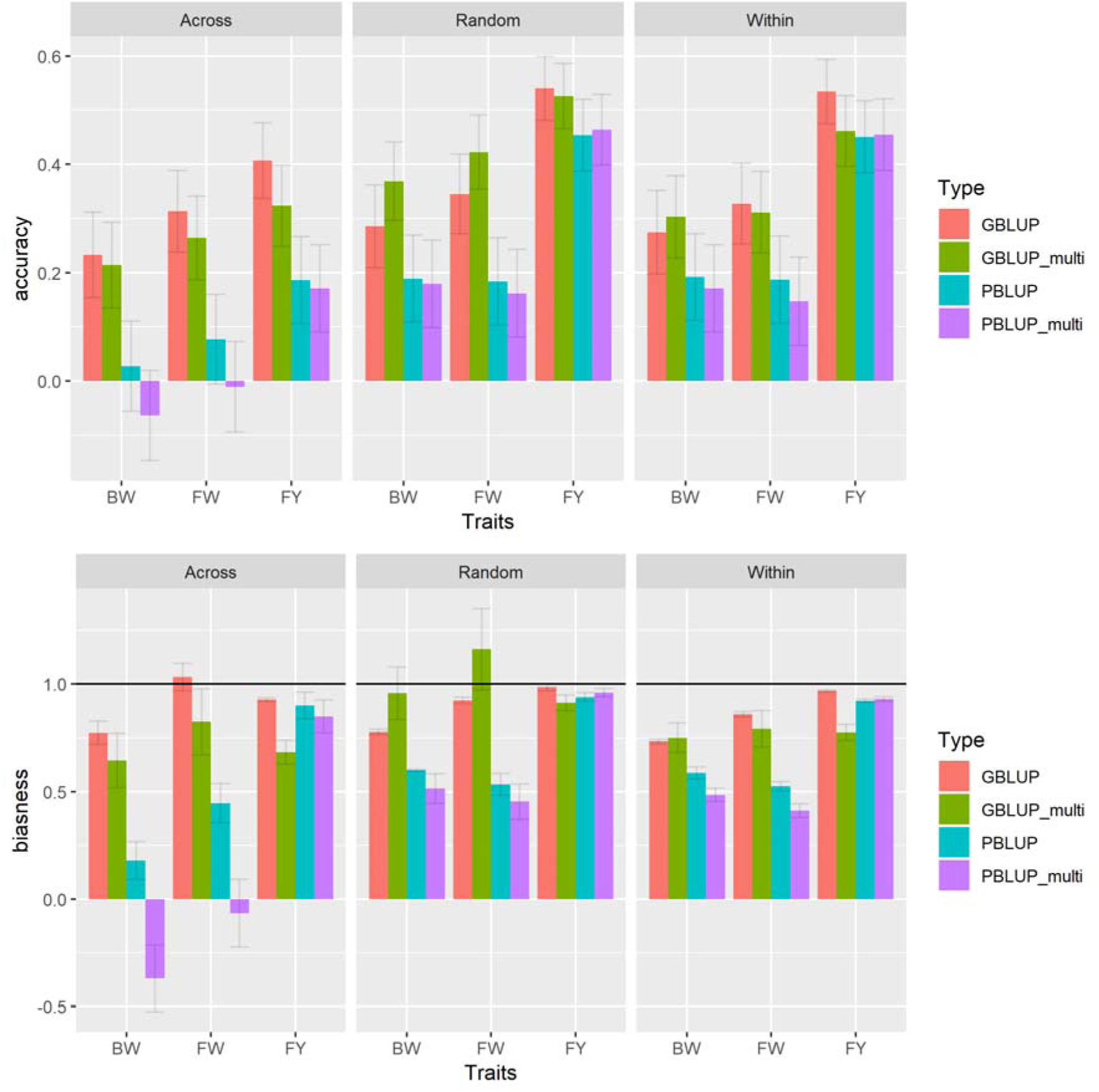
First figure showing accuracy of prediction and second one showing prediction bias. “Across family cross-validation” method is presented as “across”, “within family cross-validation” method as “within” and “random cross-validation” method as “random. The models with univariate approach are shown as PBLUP and GBLUP, whereas the models with multivariate approaches have suffix “multi” in the models. The lines in the bar charts represent ± standard errors.

### Prediction bias

Estimates for the prediction bias using PBLUP vs GBLUP models in different cross-validation methods are presented in Figure 3. Pedigree based models were found to inflate the estimated breeding values compared to GBLUP models. The prediction bias showed similar pattern to the prediction accuracy across all the models and methods. The PBLUP multivariate models were negatively biased for BW and FW in the across model cross-validation method.

## Discussion

Genomic heritabilities have previously been reported for the commercial traits in Nile tilapia [26,27], but these studies fail to report the predictive abilities of the genomic and pedigree based models. In another study, increase in prediction accuracies was indeed reported for Nile tilapia [28], based on univariate single-step GBLUP models. Thus, to the best of our knowledge this is the first report comparing prediction accuracy using both univariate and multivariate approaches with GBLUP models and pedigree-based models in Nile tilapia. Thereby, these are the first reports on heritabilities and correlations using multivariate genomic models.

### Genomic selection increases prediction accuracy in Nile tilapia

The increase in the prediction accuracy using GBLUP models, is due to the more accurate construction of the relationship matrices with better estimation of the Mendelian sampling effects using genomics (Figure 4). Using PBLUP models all full-sibs (without own phenotype) have identical EBVs, which is the parental average. Whereas, GBLUP can capture the Mendelian segregation among the full-sibs and the putatively best (unphenotyped) candidates within a full-sib family can be identified. This explains the very low accuracy (near to 0) in across-family cross-validation methods using PBLUP. Thus, the benefit of using genomics to predict the breeding values is very significant for invasive traits, where the breeding values of the animals in different full-sib families might have to be predicted based on phenotypes on other full-sib families. For example, disease challenge test in a handful of full-sib families due to expensive phenotype measurement.

**Figure 4:**
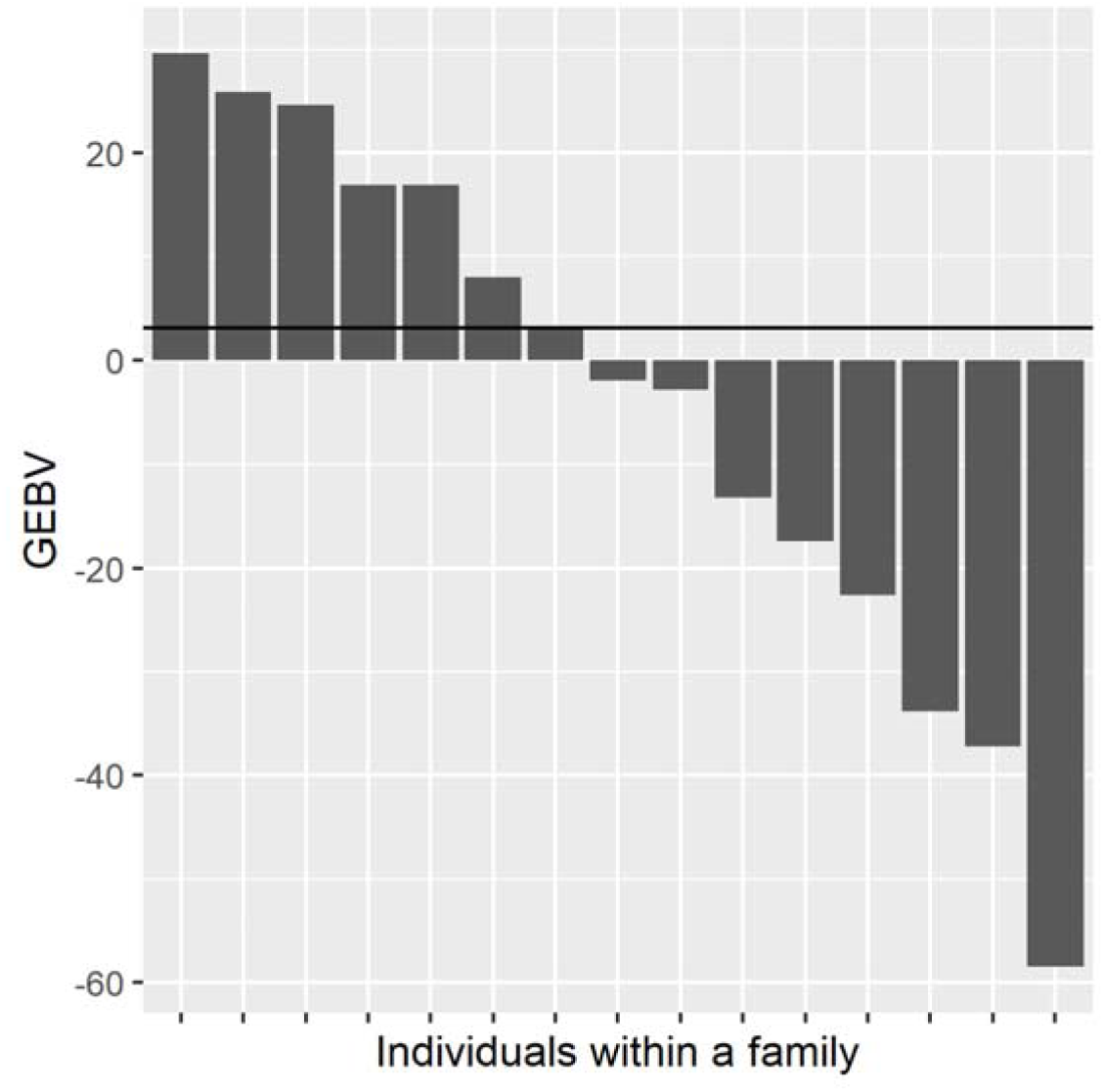
Distribution of GEBVs (g) for BW in a family with 15 offspring. The cross-validation using PBLUP predicted only one breeding value (shown as a horizontal line) for all the full-sibs. Whereas the GBLUP predicted different breeding values for all the full-sibs based on Mendelian segregation.

The lower prediction accuracy for the traits BW and FW, compared to FY across all the models and approaches may be related to the heritability and the genetic architecture of the trait [29,30]. The expected accuracy of prediction has been given as [31,32]:

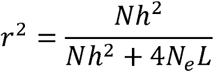

where, r is the accuracy of prediction, N is the number of animals in the training set, h^2^ is the heritability, N_e_ is the effective population size and L is the genome size in Morgan. Given the same training set and phenotypes being measured in the same animals, accuracy of prediction decreases with the decrease in heritability [33]. Joshi et al. [19,34] have shown the substantial contribution of non-additive genetic effects and maternal effects for BW and substantial contribution of maternal effects for FW in Nile tilapia. Thus, for BW and FW, maternal effect and non-additive genetic effects are part of the genetic architecture, but the model used cannot separate these effects in our data, which may affect predictive ability. Whereas, the trait FY was shown to favor simple additive model, like the model we have used in this study. Hence using the model corresponding to its genetic architecture might have increased the prediction accuracy for FY, compared to BW and FW. The mating design used in the current study made it impossible to fit complicated models to separate non-additive and maternal effects.

Further, the prediction accuracy for these commercial traits is somewhat lower than that have been reported in Nile tilapia [28] and other species [5,6]. One of the reasons for this might be our data structure. In the study we have 20 full-sib families with only one observation per family and a few more families with only 2 or 3 animals per family. Prediction of the phenotypes for the individuals in these families based on the information from other families gives lower accuracy, which might have affected our overall value of the prediction accuracy. Another reason for overall lower prediction accuracy might be the sample size. It has been stated that 2NeL number of animals are required to achieve accuracies higher than 0.88 [33], and the accuracy decreases with the decrease in the sample size and vice versa. In GST® strain of Nile tilapia, this suggests that we need at least 2304 animals (Ne= 83 (unpublished result) and L= 14.70 [14]) in training set for higher prediction accuracy, but this study uses 1444 samples.

### Multivariate approaches were not found to increase the prediction accuracy in Nile tilapia

Multivariate approaches account for the genetic and phenotypic correlation between the traits and are supposed to increase the prediction accuracy and decrease the bias [35] depending on the genetic correlation between the traits. On one hand, various studies have shown an increase in prediction accuracy for traits with low heritabilities, when used together with a correlated trait of higher heritability [36]. On the other hand, it has also been shown that when the genetic correlation between the traits is low (like BW and FY in our case), there is no improvement in accuracy using multivariate approaches over univariate approaches [37,38]. No consistent differences in the prediction accuracy was found between univariate and multivariate GBLUP models which might also be related to the types of traits used in this study. The three traits studied are not independent, as FW is a part of BW, while FY is a ratio of the two former traits.

The obvious question now is; which method is the best and should be used in the evaluation in the current Nile tilapia breeding program. Theoretically, the models giving best prediction value, minimising mean-squared error and giving unbiased estimates of the EBVs should be used [42,43], whereas practically this also depends on the selection schemes, for example the selection among the single generation of individuals, like in Nile tilapia, depends only in the prediction accuracy, as they share the common mean and bias is not concern. Whereas, it is strongly recommended to consider bias in the selection of the prediction model, if the aim is to compare between multiple generations and to predict the genetic potential of the young animals [39].

### Estimates of variance components and heritabilities

Our study showed moderate heritabilities for BW, FW and FY, which have also been reported in previous studies [19,26,28,34,44–46]. Similarly, the genetic and phenotypic correlations between the traits are similar to what has been published earlier [47], but there are some studies indicating a positive genetic correlation between BW and FY [44,46], while our estimates are negative. Negative genetic correlation between BW and FY suggests a relatively larger increase in head, gut and/or skeleton tissues with increasing body size, which is undesirable. Few studies recognize that the variance parameters and the corresponding heritabilities obtained using different relationship matrices, for example numerator and genomic relationship matrices in PBLUP and GBLUP models in our study, are different estimates for different base population. Hence, re-scaling of the relationship matrices to the same base population [25] is necessary to make sense of the comparison as it has been shown that the large differences in the pedigree and genomic based heritabilities can be accounted for by this difference [25,34,48]. Hence, it will not be wise to compare our estimates of heritabilities with the published estimates without converting them to the same base (these kinds of estimates are difficult to come by for Nile tilapia).

The difference in heritabilities using PBLUP and GBLUP models were high in the univariate approach compared to the multivariate approach. Comparing the heritabilities based on different approaches, FY gave similar heritabilities for both multivariate and univariate approaches, given the same model. For BW and FW, multivariate models gave slightly higher (but not significantly different) heritabilities compared to univariate models, whereas PBLUP models gave generally higher heritabilities compared to GBLUP models. This suggests that the markers used in GBLUP was not able to capture all genetic variance (especially if the family structure is not that strong).

An earlier study [34] has also shown the higher pedigree based heritabilities compared to genomics for these three traits (which were scaled to the same base) for the population out-crossed from generation 22 of the GST® strain (in this study we are using generation 27 of the GST® strain). Comparing the value of the estimates, heritabilities obtained using GBLUP models in our study were similar to theirs, whereas the heritabilities using PBLUP in our study was lower than theirs.

## CONCLUSION

Genomic selection is beneficial to the Nile tilapia breeding program as it increases prediction accuracy and gives more unbiased estimates of the breeding values compared to the pedigree. It is recommended to use an univariate GBLUP approach in the routine genetic evaluation for the commercial traits in Nile tilapia.

## List of abbreviations

Acronym: **Full Form**
BW: Body Weight at Harvest
FW: Fillet Weight
FY: Fillet Yield
GBLUP: Genomic Best Linear Unbiased Prediction
GST: GenoMar Supreme Tilapia
G(EBVs): (Genomic) Estimated Breeding Values
PBLUP: Pedigree Best Linear Unbiased Prediction

## Declarations

### Ethics approval and consent to participate

Not applicable

### Consent for publication

Not applicable

## Availability of data and material

The data used in the study are from commercial family material. This information may be made available to non-competitive interests under conditions specified in a Data Transfer Agreement. Requests to access these datasets should be directed to Alejandro Tola Alvarez.

## Authors’ contributions

RJ did the statistical analysis and wrote the initial draft of the paper, AS contributed to the draft and was responsible for genotyping and microsatellite-based pedigree construction, MDV supervised the experiments, phenotyping and collection of fin samples in the farm, ATA conceived the study, JØ supported in the statistical analysis and all authors contributed to the discussion of the results and writing of the final version of the paper.

## Authors’ information (optional)

Not applicable

## Supplementary

**Figure S1:**
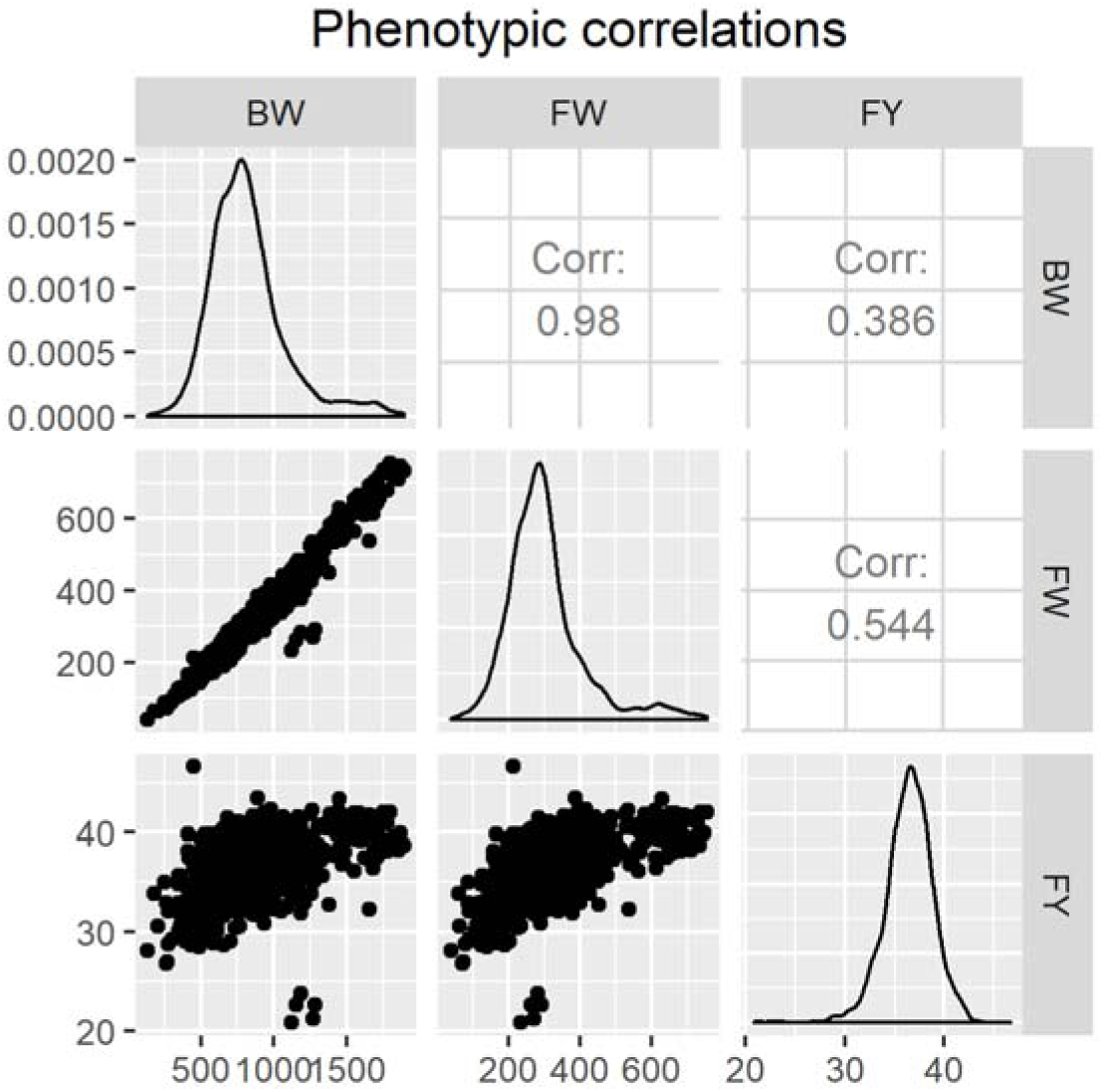
Scatterplots and correlation between different phenotypes. Phenotypic correlation between the traits is not corrected for fixed effects in the plot. Table 3 shows the phenotypic correlation corrected for fixed effects.

